# piCRISPR: Physically Informed Deep Learning Models for CRISPR/Cas9 Off-Target Cleavage Prediction

**DOI:** 10.1101/2021.11.16.468799

**Authors:** Florian Störtz, Jeffrey Mak, Peter Minary

## Abstract

CRISPR/Cas programmable nuclease systems have become ubiquitous in the field of gene editing. With progressing development, applications in *in vivo* therapeutic gene editing are increasingly within reach, yet limited by possible adverse side effects from unwanted edits. Recent years have thus seen continuous development of off-target prediction algorithms trained on *in vitro* cleavage assay data gained from immortalised cell lines. It has been shown that in contrast to experimental epigenetic features, computed physically informed features are so far underutilised despite bearing considerably larger correlation with cleavage activity. Here, we implement state-of-the-art deep learning algorithms and feature encodings for off-target prediction with emphasis on *physically informed* features that capture the biological environment of the cleavage site, hence terming our approach piCRISPR. Features were gained from the large, diverse crisprSQL off-target cleavage dataset. We find that our best-performing models highlight the importance of sequence context and chromatin accessibility for cleavage prediction and compare favourably with literature standard prediction performance. We further show that our novel, environmentally sensitive features are crucial to accurate prediction on sequence-identical locus pairs, making them highly relevant for clinical guide design. The source code and trained models can be found ready to use at github.com/florianst/picrispr.

## Introduction

The clustered regularly interspaced short palindromic repeats (CRISPR) sequence family was first described in *E. coli* in 1987 [1], but it took until 2007 to recognise it as a part of the viral defense system of most archaea and bacteria [2]. Exogenous viral DNA is cleaved off by specialised nuclease enzymes, coded for on genomic regions which are often adjacent to CRISPR and hence named CRISPR-associated (Cas). Cleaved-off regions are subsequently incorporated into the CRISPR sequences, which act as a viral history of the respective cell, stabilised by the palindromic nature of their saved states which results in stable secondary structures [3]. From there they can be transcribed to crRNA and invading copies of them can subsequently be rendered inactive. Researchers have used this ability for programmable genome editing in many eukaryotic species, complementing strategies such as zinc-finger nucleases (ZFNs, [4]) and transcription activator-like effector nucleases (TALENs, [5]).

We concentrate on the effects of the wild-type Cas9 protein gained from *Staphylococcus pyogenes*. The crRNA which is originally responsible for recognition of a 20bp viral sequence forms an active complex with the tracrRNA, called single guide RNA (sgRNA), of about 50bp length [6]. Homology of the crRNA part with a 20bp region in the genome results in annealing of the sgRNA with one strand of this region, which we call ‘target strand’. Binding happens when the interaction of the 3bp protospacer-adjacent motif (PAM) on the opposite, non-target strand with the Cas9 protein is favourable [7]. For *S. pyogenes* Cas9, this is the case for an ‘NGG’ PAM where N stands for an arbitrary base (A, T, C, G).

Tertiary DNA structure, such as nucleosome octamers, can occlude or expose different regions of DNA and hinder Cas9 access [8]. After binding has taken place, nuclease-active enzymes within Cas9 can cleave the double-stranded DNA 3bp upstream of the PAM. Due to the stochastic, energy-driven nature of both the binding and the cleavage process, we expect a distribution of cuts over the whole genome, including undesired off-target effects which could possibly have catastrophic consequences, such as knocking out tumor suppressor genes like p53 and Rb [9].

We noticed that repositories of off-target cleavage data contain a significant amount of data points which match in both guide and (off-)target sequence and differ only in the biological environment of the respective loci (see Figure 1). Capturing this environment is therefore instrumental in providing accurate predictions of cleavage activity. We recently found that computed nucleosome organisation-related features correlate better with cleavage frequency values than experimental epigenetic markers (DNase I, RRBS, CTCF, H3K4me3) [10] which have heretofore been the literature standard for cleavage prediction models. These computed features also surpassed epigenetic markers in terms of their feature importance in preliminary cleavage activity prediction models which had access to both computed and experimental epigenetic features. We therefore aim to make full use of this novel class of features by embedding them in a rich feature set, including DNA/RNA sequence and context sequence-based features.

**Figure 1:**
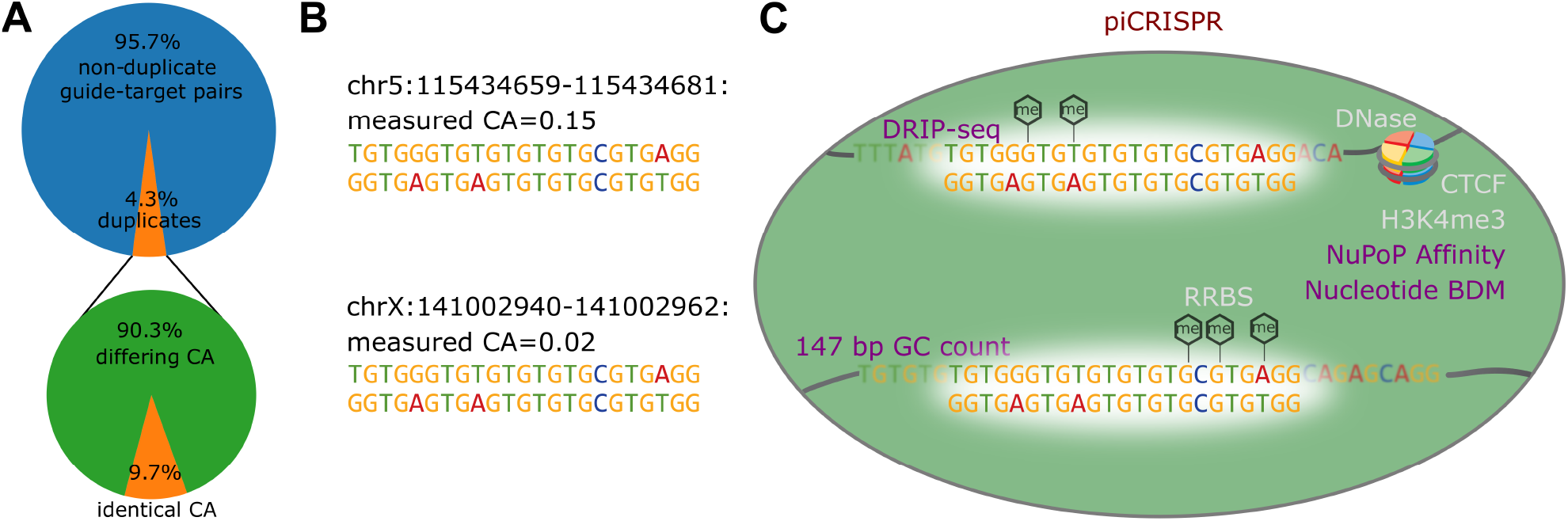
**A** The crisprSQL dataset contains an appreciable amount of perfect guide-target duplicates. We only consider data gained from human cell lines and putative off-targets which we generated based on sequence similarity. 8,922 of 230,274 data points have at least one guide-target duplicate within this set which differs in cleavage activity (CA). As the example from our dataset in panel **B** shows, such a pair looks identical to purely sequence-based prediction algorithms. They might therefore not predict dangerous off-target effects. **C** piCRISPR remedies this by taking into account the biological environment of the cleavage site based on a range of features beyond guide and (off-)target sequence. So far, prediction algorithms have used features related to chromatin organisation (CTCF, [20]), chromatin accessibility (DNase-Seq), DNA methylation (RRBS) and histone methylation (H3K4me3, [21]). We on the other hand use features pertaining to the 147 bp sequence context around each (off-)target nucleotide: GC count, sequence complexity (BDM, [19]) and nucleosome positioning information (NuPoP, [8]) which introduce unprecedented sensitivity to the biological environment of the cleavage site. Using these, piCRISPR can correctly rank the two example loci given here.

With a considerable amount of cleavage prediction algorithms present in literature [11, 12, 13, 14, 15], we present here a choice of two model architectures, two encodings and two sets of features, yielding a total of eight combinations. We scrutinise these according to both prediction performance and interpretability. Besides improving prediction accuracy and capturing off-target effects that might so far have gone unnoticed, this will also generate insight into the biological environment that influences CRISPR cleavage.

## Methods

### Data Source

In order to achieve maximum transparency and comparability, we use guide-target pairs from the crisprSQL dataset [16], curated by our group. It is a collection of 17 base-pair resolved off-target cleavage studies on Sp-Cas9, comprising 25,632 data points and is larger than most datasets used to train prediction algorithms to date. It contains data on various cell lines, mainly U2OS, HEK293 and K562. We have chosen to use version 26/05/2020 of the database which does not include T-cell data from [17] in order to avoid introducing a considerable cell line imbalance. Furthermore, the evaluation of our modelling on on-target datasets is beyond the scope of this work due to their different underlying experimental techniques and cleavage quantification measures.

Experimental data points containing guide and target loci, sequence, cell line, assay type and cleavage frequency have been completed and enriched by sequence context as well as the `CRISPRoff` score, an empirical estimate of the (off-)target binding energy calculated in [18].

Besides these established features, we propose the usage of nucleosome organisation-related features [10] which add an unprecedented level of sensitivity towards the biological environment of the cleavage site (see Figure 1C). In this publication, we trained a preliminary cleavage prediction model on 13 distinct nucleosome organisation-related scores all based on the 147 bp context around each (off-)target nucleotide (see Supplementary Material) as well as the four literature-standard epigenetic markers named above and include the three scores of highest feature importance: GC count, Nucleotide BDM [19] and NuPoP Affinity [8].

GC count refers to the relative proportion of G and C nucleotides within the 147 base pair window around a given target nucleotide. Nucleotide BDM refers to a training-free method that approximates the algorithmic complexity of a DNA sequence. Low values of Nucleotide BDM have been shown to correlate with proximity to nucleosome dyad positions [19]. NuPoP refers to a duration Hidden Markov Model trained to predict the base-pair specific nucleosome affinity of a given (off-)target sequence. For the precise calculation of these three features, we refer to [10].

### Data Augmentation

In order to increase the size of the training set, we extend it by those putative off-target sites along the respective genome which had fewer than seven mismatches to each respective guide sequence, omitting the (off-)target locus itself. It was ensured that the protospacer adjacent motif (PAM) at the very end of the guide sequence was either the canonical 5’-NGG-3’ characteristic of SpCas9 [22], or one of the noncanonical forms 5’-NGA-3’ and 5’-NAG-3’ observed in [23]. If a genome-wide off-target detection method has not detected cleavage at a locus within the genome that satisfies these criteria, we deem the cleavage activity at this point to be zero. This yielded 310,142 total guide-target pairs, making the complete data set highly imbalanced. Sticking with the convention in literature, we refer to this process of extending the number of data points as *data augmentation*. For this work, we concentrated on the 251,854 data points originating from a human cell line or synthetic human DNA.

#### Labels

For classification, we define the negative class as all data points with cleavage activity (CA) values below the lowest reported assay accuracy of 10^−5^, combined with the set of putative off-targets. In order to achieve comparability between different studies for regression tasks, we perform a nonlinear Box–Cox transformation [24] to transform the cleavage rates to approximate a Gaussian with zero mean and variance *σ*^2^ = 2, similar to the approach in [25] and [13]. Cleavage activity values below the lowest reported assay accuracy of 10^−5^ as well as putative off-targets were set to − 2*σ*^2^ = − 4, and transformed values clipped to the [− 4, 4] range. This is an empirical choice based on the shape of the resulting distributions.

### Feature Encoding

We employ two different feature encoding schemes which occupy different points in the tradeoff between sparsity and interpretability. The first was introduced in [13] and consists of one-hot encoded representations of the guide and target sequence which have been combined using a bitwise OR operation. In order to make up for the loss of information that this operation causes in terms of mismatches, two additional channels are added containing information about the directionality of the bases involved in the mismatch, i.e. which of the two entries describes the target and which the guide nucleotide. Note that the guide nucleotide is first translated into its corresponding target protospacer nucleotide. We call this encoding the 6 × 23 encoding based on the resulting shape of the sequence matrix (see Figure 2A).

**Figure 2:**
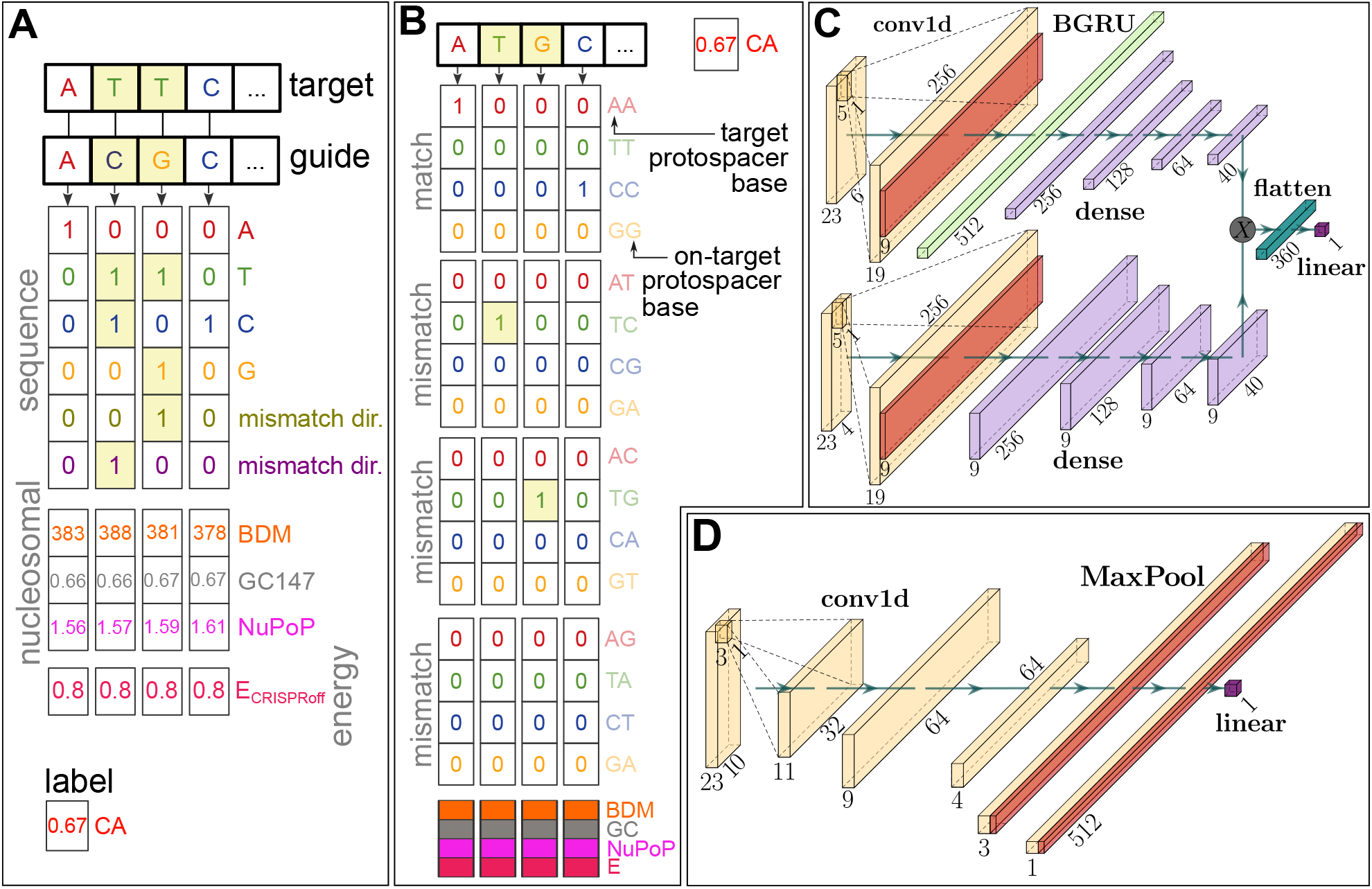
Overview of our two feature encodings and model architectures. **A** 6 × 23 encoding: (Off-)target and guide (on-target protospacer) sequences are one-hot encoded and copied together using a bitwise OR operation. In order to make this encoding lossless, two channels are added that encode which nucleotide is on the guide and which on the target, respectively (in the case of a mismatched interface). This is identical to the encoding used in [13]. The nucleosome positioning channels (147 bp GC count, Nucleotide BDM [19], NuPoP Affinity [8]) of the target as well as the `CRISPRoff` free energy estimate score [27, 18] are concatenated to this matrix. The Box–Cox transformed cleavage activity (CA) is used as a label. **B** The 16 × 23 encoding uses a 16-letter alphabet which explicitly contains information about the precise nature of the mismatch. **C** Bidirectional gated recurrent unit architecture as used in our RNN model, modelled after the network in [28]. Upper and lower arm of the network contain the sequence and nucleosomal/energy information, respectively. **D** Convolutional neural network architecture used in our CNN model, comparable with the model in [11]. Dimensions in both model architectures are valid for the 6 × 23 encoding (panel A).

Based on the energy-driven nature of binding and cleavage, we hypothesise that mismatched interfaces affect binding in a totally different way than matched interfaces. This has so far not been recognised in detail by off-target prediction algorithms. Since the 6 × 23 encoding contains the information about the precise nature of a given interface only implicitly, we decided to include a further encoding which does so explicitly. This uses a one-hot representation using the 16 letter cross product between guide and (off-)target nucleotide, and is hence termed the 16 × 23 encoding (see Figure 2B). This is similar to the encoding scheme in [26].

Both matrices are then concatenated with a matrix of base-pair resolved nucleosomal features and the `CRISPRoff` value of the given target-guide interface repeated along the sequence axis. Exploring latent representations of guide or target is not within the scope of this work, given that it further complicates comparison between models.

### Model Architectures

Literature contains a wealth of model architectures commonly used to predict CRISPR cleavage. Currently, successful model architectures for learning-based cleavage prediction fall in one of three categories: tree-based methods [25], convolutional neural networks (CNN, [11, 12]) and recurrent neural networks (RNN, [13, 28, 15]). We take successful CNN and RNN architectures present in the field and adapt them to the task of off-target prediction using various encodings of the features described above.

Our CNN model is comparable to the architecture described in [11]. There, the outputs of two separate, convolutional layer-based encoders for guide and (off-)target are concatenated channel-wise (forming the Siamese part of the network) and serve as input for a convolutional classifier (the conjoined part). Since both encodings scrutinised in this publication combine guide and target sequences, we only utilise one arm of this Siamese network (see Figure 2D). We have made various adjustments to this architecture based on training stability and validation set performance (see the Supplementary Text).

Our RNN architecture is modelled after the bidirectional gated recurrent unit (BGRU) on-target prediction model from [28]. Here, a BGRU layer is used to make use of the relevant longer-range dependencies between sequence features that would go unnoticed by a CNN of manageable kernel size. In order to make this type of architecture usable for off-target prediction, we feed a combination of guide and target sequence as described above into the sequence arm of the network, and the nucleosomal and energy features in a separate arm (see Figure 2C). Layer dimensions in both arms were adjusted to the shapes of their respective inputs.

### Model Training & Evaluation

Given the imbalance of validated/measured and non-validated/augmented data points, we employ a bootstrapping strategy as suggested in [29], where training batches on average contain equal numbers of both classes. For regression (classification), early stopping is based on the mean squared error (binary cross-entropy) loss on half of the test set, where the other half is reserved for evaluation.

The CNN models are trained in the same way, with hyperparameters of `batchnorm_momentum`=0.01, Gaussian noise with *μ* = 0, *σ* = 0.01 and Adam learning rate 10^−3^.

The RNN models are trained for 100 epochs, where batches of 10,000 points are sampled each epoch out of a class-balanced sample of 50,000 data points. We replicate the transfer learning approach taken in [28] with adjustments to increase training stability and generalisation performance as detailed in the Supplementary Text. Dropout probability was 0.2 and the Adam learning rate was 10^−3^.

#### Testing Scenario 1: held out studies

In this scenario we hold out studies [30, 31, 32] from the training set. These studies have not been included in the training set for the state-of-the-art off-target prediction algorithm CRISPR-Net [13], such that they remain an independent test set to compare CRISPR-Net and piCRISPR side by side. The inherent class imbalance in this test set is 1:103.96. 22 % of the unique guides within the training set have at least one corresponding guide in the test set with five or fewer mismatches, indicating a satisfactory independence between training and test set.

#### Testing Scenario 2: literature comparison

In this scenario we use the CIRCLE-seq [33] dataset as the held out test set, as was done in [15]. The exact test set has been replicated using the code provided by the authors, such that comparison values could be taken straight from publication [15]. Nucleosomal and empirical energy data was filled in using the crisprSQL dataset.

#### Testing Scenario 3: set of duplicate pairs

In this scenario, we scrutinise our hypothesis that an environmentally sensitive feature set is fit to not only increase prediction performance overall, but especially for given groups of identical guide-target sequence pairs. To this end we calculate two quantities: First, the mean squared error (MSE) between the predicted regression scores and the ground truth cleavage frequencies within each of the 2,703 groups. Second, the average proportion of the true cleavage activity difference for two points within a given group which the model predicts. This is zero for purely sequence-based models and unity for an ideal predictor. This quantifies how faithful a model is to the differences in biological environment for a given pair. In order to emphasise small deviations which preserve the rank of predicted cleavage activities, we use the cubic root as a sign-preserving nonlinearity and term this quantity *relative difference*. We consider the resulting distributions of both of these quantities for different feature sets.

### Model Explanation

We obtain feature importances using the model-agnostic Shapley value explainer library SHAP [34]. Since piCRISPR wraps the feature encoding inside a given model, we retain full explainability of input features even for non-invertible encodings. In this way, using the two encodings detailed above, we obtain an unprecedented, context-sensitive resolution of sequence-based features.

Sticking with the convention set by the SHAP library [34], we calculate global SHAP values as the mean of the absolute value of SHAP values across data points in the explanation set, which is a random subset of 500 points from the held out test set. In order to show not only the magnitude but the direction in which a given feature influences the model’s prediction, we multiply each feature’s global SHAP value by the sign of the average SHAP value of all data points whose value is larger than the median of that feature.

### Command line usage of our models

We have implemented a command line interface with which piCRISPR predictions can readily be obtained. For maximum usability, the model automatically uses default feature values in case a certain feature was not provided, thereby lowering prediction performance (see Figure S4). The default value of a given feature is defined as the average feature value of the set of those crisprSQL data points which lie within a 20% interval around the mean SHAP value. This means that high-accuracy piCRISPR predictions can be obtained in a user-friendly way, even when providing only guide and (off-)target sequence. Our online repository contains hands-on examples on this.

## Results & Discussion

### Testing Scenario 1

Figure 3 shows the regression and classification performance of our piCRISPR-implemented models, with the 6 × 23 RNN model yielding the highest benchmarks. As mentioned in [29], the area under precision-recall curve (AU-PRC) is a much more suitable measure than the area under receiver operating curve (AU-ROC) for off-target prediction, since in clinical application, false negatives have far more adverse effects than false positives. The addition of the nucleosomal features considerably improve model performance according to all benchmarks, supporting our hypothesis that nucleosomal features can serve as a key ingredient to cleavage prediction.

**Figure 3:**
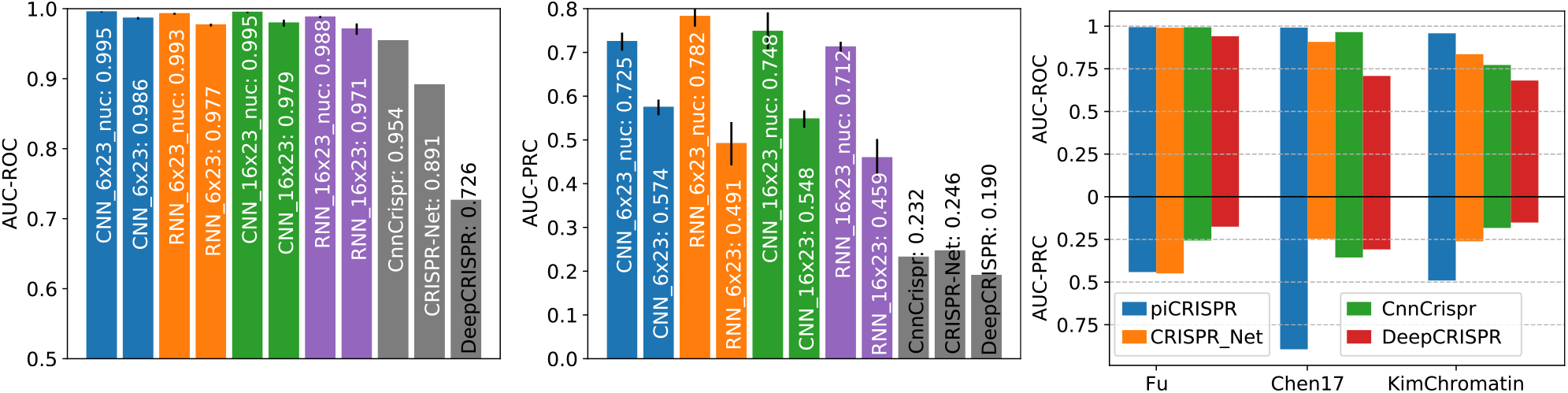
Comparison of piCRISPR models with published algorithms. All models were tested on held out studies [30, 31, 32] (testing scenario 1). Non-validated data points have been oversampled in the test set to match the class imbalance of 1:79.35 found in the dataset I-1 from [13]. piCRISPR models have been trained on the remaining data points within the crisprSQL data set. **Left two panels**: Comparison with three published off-target prediction algorithms [11, 13, 26] that were run on this test set. Within a model family of the same colour, the model labelled “nuc” contains nucleosomal features whereas the other does not. piCRISPR training and testing have been repeated 5 times to obtain mean and standard deviation as shown. For the underlying ROC and PRC curves see Figure S1. **Right panel**: AUC-ROC and AUC-PRC benchmarks for the RNN 6 × 23 model with nucleosomal features, resolved by individual study within the held out test set.

A direct comparison with prediction results obtained from the published versions of CnnCrispr, CRISPR-Net and DeepCRISPR on the identical held out test set shows that piCRISPR achieves higher classification benchmarks in terms of areas under ROC and PRC curve for all three individual studies contained in the test set, except for study [30] for which piCRISPR and CRISPR-Net achieve comparable benchmarks.

### Testing Scenario 2

Figure 6 shows that when testing on the CIRCLE-seq dataset [33], piCRISPR performance drops slightly as compared to testing scenario 1. Especially RNN models generalise slightly worse to this study. We still observe that nucleosomal features enhance the performance of a model, and piCRISPR still outperforms both CRISPR-IP and CRISPR-Net models according to area under PRC curve.

### Testing Scenario 3

Table 1 shows that the model performance, measured by the mean squared error of predictions within a group of data points that share both guide and target sequence, is considerably improved by introducing features beyond sequence information (left column). The resulting distribution of MSEs is shown in Figure S9.

**Table 1:**
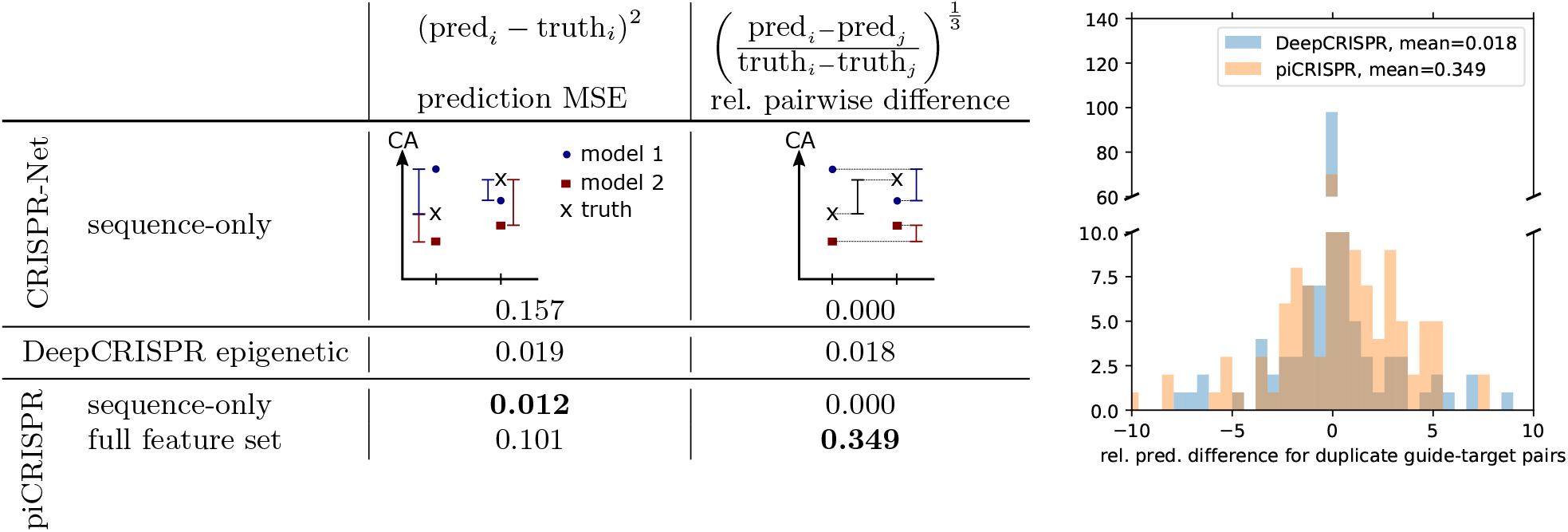
Benchmark quantities gained on the subset of duplicate guide-target sequence pairs (testing scenario 3) using our 6 × 23 CNN model as well as the CRISPR-Net [13] and DeepCRISPR [11] models for comparison. For piCRISPR, we give a sequence-only version of the model in which nucleosome organisation related features and the empirical energy estimate have been set to a default value across all data points. **Left column**: mean squared error (MSE) between predicted cleavage score and ground truth cleavage activity, averaged over all groups of identical guide-target sequence pairs. **Right column**: How faithful a model is to the differences in biological environment for a given pair within such a group is measured by the average proportion of the true cleavage activity difference which the model predicts. This is zero for purely sequence-based models and unity for an ideal predictor. To emphasise small deviations which preserve the rank of predicted cleavage activities, we use the cubic root as a sign-preserving nonlinearity and term this quantity *relative difference*. **Right panel**: Example distributions of relative pairwise difference for the two models. All underlying distributions are shown in Figure S9.

Looking at the relative pairwise difference, we observe that introducing features beyond sequence leads to an increase of the average proportion of true cleavage frequency differences between points of differing biological environment which is captured by the model. This is true for both DeepCRISPR and piCRISPR. Whilst the full feature set in piCRISPR achieves the highest proportion in comparison, it is the piCRISPR sequence-only model that achieves the lowest overall mean squared error. This indicates that a low mean squared error does not necessarily go hand in hand with the model drawing the correct conclusions from environmentally sensitive features. This can be seen as well when considering the comparably small relative pairwise difference that is recovered by the DeepCRISPR model from the literature-standard epigenetics channels to which it has access.

### Feature importance

Due to its comparatively stronger prediction benchmarks between testing scenarios 1 and 2, we use the 16×23 CNN classification model in testing scenario 1 to extract feature importance values of unprecedented resolution. Figure 4 shows that the model draws on sequence features which stem from mismatched interfaces differently than on those from matched interfaces, supporting our hypothesis that this differentiation is not only physically indicated but also backed by the model’s behaviour. Global SHAP values suggest that the preference of the variable PAM nucleotide at position 21 is contingent on the specific sgRNA–DNA interface formed. We recover the preference for cytosine at position 17 [11, 35] as well as position 20 [11, 28] found in literature for matched interfaces. However, for mismatched interfaces, cytosine is disfavoured. Whilst we cannot recover a strong preference for the variable PAM nucleotide at position 21 for matched interfaces, we observe the preference for guanine reported in literature [11, 28, 35] for mismatched interfaces. This supports the notion that a concentration on guide-target interfaces rather than pure base identities is necessary for off-target prediction, and that deeper insight is required than the notion of a preferred base at a specific position. It therefore appears necessary to consider mismatch interfaces together with sequences in the desired genome, not just the sequence of the putative guide, for sgRNA design.

**Figure 4:**
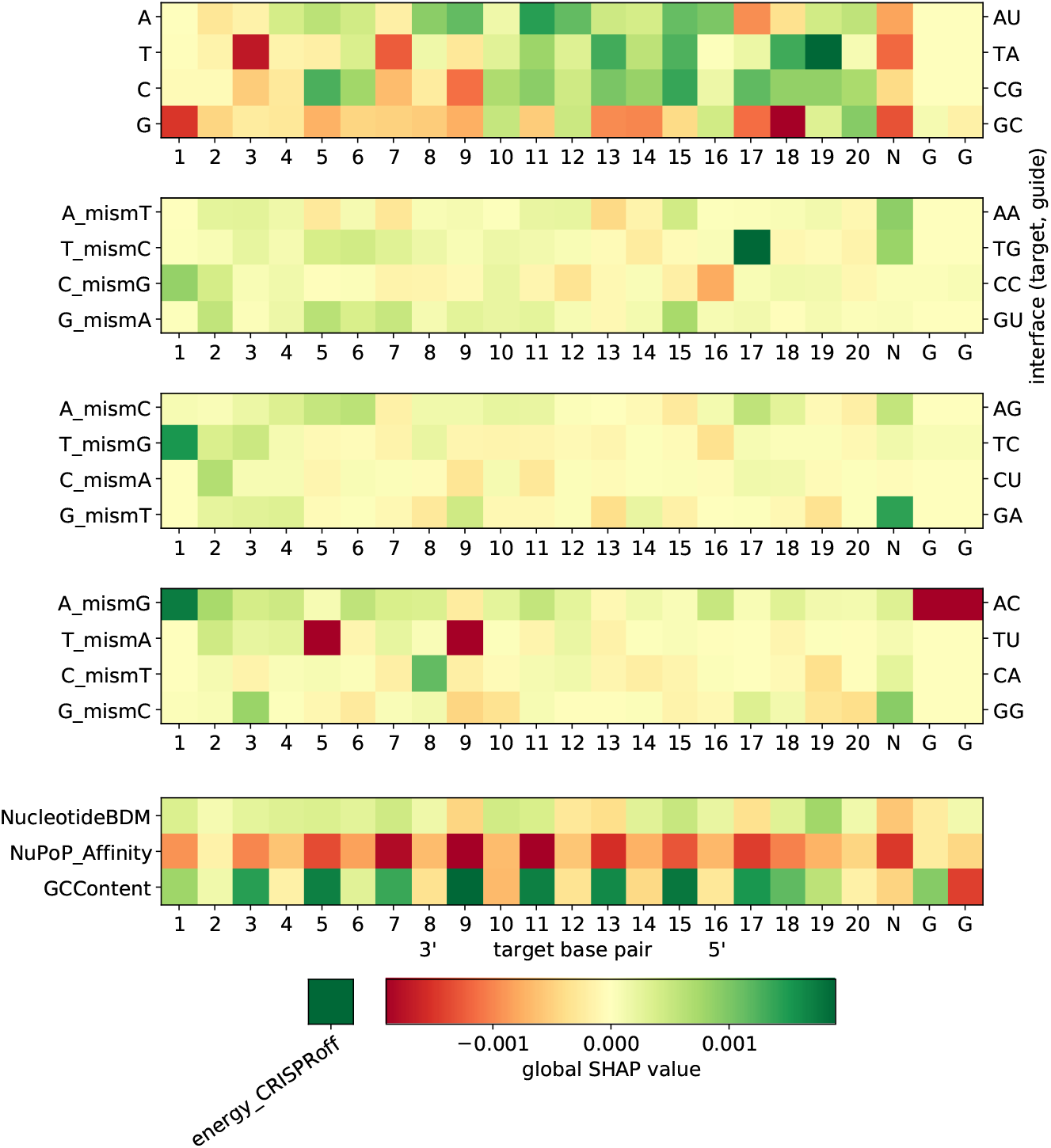
Global SHAP values for the 16 × 23 CNN classification model. Negative global SHAP values (red) indicate an average predicted decrease in guide activity for the respective feature. Mismatch channels (middle three heatmaps) can be represented by the (off-)target and on-target protospacer nucleotides (left vertical axis) as well as the physical base pair interfaces (right vertical axis), such that `A_mismT` describes all configurations in which an adenine on the target strand faces an adenine on the sgRNA. The bottom heatmap visualises the influence of our chosen set of nucleosomal organisation features on cleavage activity. A bar representation of this can be found in Figure S5.

**Figure 5:**
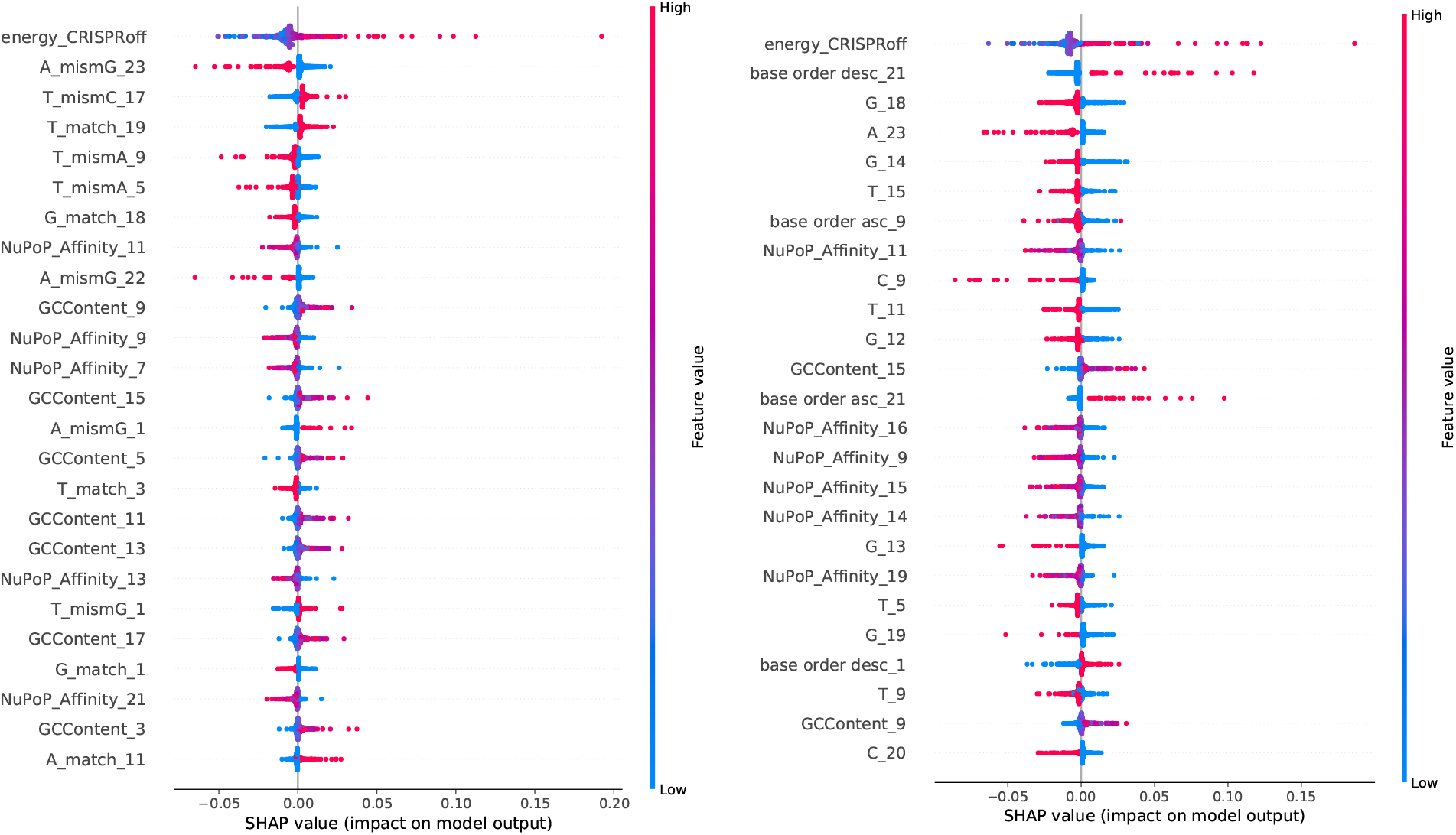
Base-pair resolved global SHAP values for the 16 × 23 (left panel, see Figure 4) and 6 × 23 (right panel, see Figure S7) CNN classification models. SHAP values have been obtained on the held out studies [30, 31, 32] from the crisprSQL dataset. Note that high values of the NuPoP Affinity feature (red dots), i.e. highly positioned nucleosomes, always influence the model towards reduced cleavage activity (negative SHAP value).

**Figure 6:**
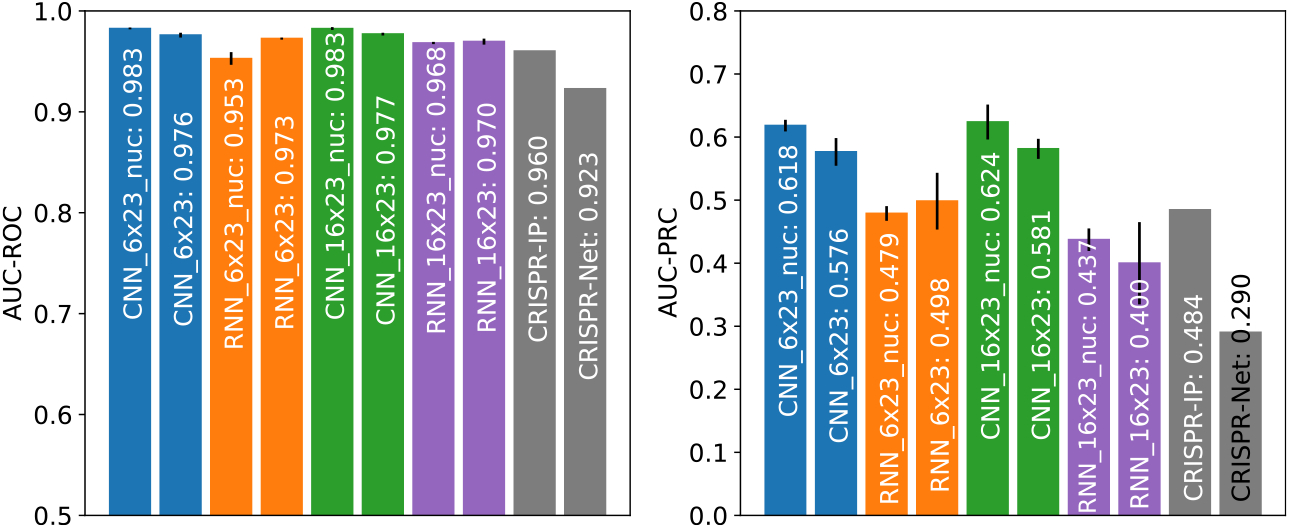
Comparison between piCRISPR, CRISPR-Net [13] and CRISPR-IP [15], with comparison values for the latter two models taken from [15]. All models were tested on a the subset of the CIRCLE-seq dataset [33] as given in [15] (testing scenario 2). Note the slightly reduced performance of the RNN models compared to testing scenario 1 (Figure 3).

Note that due to the low prevalence of non-NGG PAMs in our dataset, as has been our choice when augmenting it with putative off-targets, the model attributes little importance to the two 5’ GG base pairs. We observe the blind spot of mismatch discrimination by the REC3 domain of Cas9 around nucleotide 7 (see also Figure S5) which has been reported in a recent cryo-EM structural study [36] and results in reduced importance of sequence features pertaining mismatched interfaces in this region. At nucleotides 3–5 and 9–11, where mismatch detection by the REC3 domain of Cas9 is high, we observe a mismatch-induced reduction in cleavage activity. We further observe a PAM-distal ‘preference zone’ and a PAM-proximal ‘avoiding zone’ of mismatches when averaging over feature importance values by nucleotide, which has been observed in computational [11] as well as cryo-EM [36] studies.

The model draws heavily on the empirical energy estimate feature *E*_CRISPRoff_ which yields the largest global SHAP value. We further observe a considerable correlation between its value and the SHAP value attributed to it by the model (Figure S8). An energy score of *E*_CRISPRoff_ = −1.15 has a neutral influence on cleavage activity in the 16 × 23 CNN model, with higher (lower) values yielding larger (smaller) SHAP values, i.e. a positive (negative) influence on cleavage activity.

When considering nucleosome positioning-related feature channels, we see that the 147 bp GC content around each nucleotide has a net positive influence on cleavage activity. Similar to the argument in [10], this can be attributed to the increased bendability of GC-rich DNA [37] which is beneficial to Cas9 sequence readout during binding [38]. We further observe that the NuPoP Affinity score ranks higher in terms of global SHAP value than most sequence features. The negative influence of nucleosome affinity can be explained by the reduced accessibility of high-affinity DNA regions, and is observed strongly between nucleotides 5 and 19. This effect has been observed in [10] as well. We further observe an overall negative influence of low Nucleotide BDM values on cleavage activity, supporting what has been observed in preliminary, non-sequence based models in [10].

This also demonstrates the importance of nucleosome-related features for cleavage prediction, and also supports the notion of chromatin accessibility influencing cleavage activity found in [39]. To our knowledge, this strong effect of a more than 10 bp wide sequence context on genome-wide off-target cleavage prediction has not been demonstrated yet. Hints of it have been seen only for smaller contexts and on-target efficacy prediction [40, 41]. In addition, our findings present an unprecedented example in which information in the 147 bp sequence context has considerable impact on the model.

A similar analysis for the 6 × 23 CNN model can be found in Figure S6 and for the 16 × 23 RNN model in Figure S7. Note that within the nucleosomal feature class, the RNN models attribute more importance to the Nucleotide BDM feature than the CNN models scrutinised here. This could in part explain their slight difference in performance between testing scenarios 1 and 2.

## Conclusion

Through the introduction of a new feature class and the careful adjustment of model architectures, we have identified a set of models which match the performance of state-of-the-art off-target cleavage prediction algorithms in direct comparison. All models are highly influenced by nucleosome organisation-related features such as histone binding affinity, which emphasises the importance of capturing the biological environment around the cleavage site when modelling cleavage activity. Our approach has shown that these computed physically informed features are fit to enhance the predictive power of cleavage prediction models and to replace experimental epigenetic markers in future modelling efforts. We have further provided an accessible, user-friendly command line interface that allows users of various disciplines to utilise all our models, even without providing a complete set of features. This all paves the way towards the prediction of off-target sites which would so far have gone unnoticed.

Our environmentally sensitive set of features reveals several novel, promising pathways towards further improvement of off-target cleavage prediction. Going forward, it could be fruitful to increase model complexity, e.g. using a 2D convolutional kernel to capture interaction between features of a single nucleotide. A 2D convolution kernel would be able to capture the base-pair resolved interaction between sequence and nucleosomal markers as well as between sequence k-mers. Further than this, our multimodal data could be fused at different stages, such that sequence, nucleosomal and energy features interact at different levels of representation of each other.

We further envision to replace the epigenetic information of the guide, which so far only copies the epigenetic information of the target DNA. This is clearly an unphysical choice. Given that a synthetic sgRNA does by design not carry epigenetic markers, a one-hot encoded dot-bracket representation of the sgRNA folding would be a more suitable choice to capture its informative properties.

## Funding

This research was funded in whole or in part by the Biotechnology and Biological Sciences Research Council (BBSRC) [BB/M011224/1, BB/S507593/1]. For the purpose of Open Access, the author has applied a CC BY public copyright licence to any Author Accepted Manuscript (AAM) version arising from this submission. Some of the presented results have been obtained using the University of Oxford Advanced Research Computing (ARC) facility (http://dx.doi.org/10.5281/zenodo.22558). The authors declare no conflict of interest.

## Notes

https://github.com/florianst/picrispr

### Competing Interest Statement

The authors have declared no competing interest.

### Summary of Updates

Look at guide overlap between train and test set. Concentrate on two model architectures & (lossless) encodings. Omit established epigenetic features. Provide additional (direct) literature comparisons.

