## Supplementary Material for "piCRISPR: Physically Informed Deep Learning Models for CRISPR/Cas9 Off-Target Cleavage Prediction"

### List of nucleosome positioning-related scores

We implemented an off-target cleavage prediction model that takes into account the following features, as well as the literature-standard epigenetic features DNase I, RRBS, CTCF, H3K4me3 from crisprSQL [1] (data not shown):

- W/S Scheme: predictor of rotational nucleosome positioning, available on the online database nuMap [2]
- YR Scheme: predictor of translational nucleosome positioning, available on the online database nuMap [2]
- LeNup (H3Q85C) [3]: CNN-based predictor of nucleosome positioning
- NuPoP [4]: Hidden Markov Model-based predictor of nucleosome positioning, yields histone binding affinity, nucleosome occupancy and Viterbi scores
- nuCpos [5]: Hidden Markov Model-based predictor of nucleosome positioning, building on NuPoP, yields histone binding affinity, nucleosome occupancy and Viterbi scores
- VanDerHeijden [6]: statistical mechanics-based method for predicting nucleosome positioning
- GC147: GC content of the 147 bp sequence around each nucleotide
- Nucleotide BDM: applying the block decomposition method [7] on the 147 bp sequence around each nucleotide
- Strong-Weak BDM: Nucleotide BDM starting from the sequence which has G/C replaced with S and A/T replaced with W

### Neural network architectures

As described above, our CNN model is based on the architecture described in [8]. Instead of a dropout layer, we use a Gaussian noise layer after the first convolutional layer in each encoder since we found this to improve validation set prediction benchmarks. Having batch normalisation layers only before and after the Siamese layers increased training stability without compromising training performance. The same was true for applying (leaky) ReLU activation functions after the last (first) three convolutional layers. The last three also contained a dimension-preserving max-pooling layer in between the convolution and ReLU layers. Furthermore, we found that in our case, removing the 256-filters convolutional layer before concatenation of the Siamese channels did not considerably affect training performance but reduced training time. We then adjusted the downstream network accordingly, which involved removing the penultimate convolutional layer and halving the number of filters in the remaining convolutional layers.

The RNN models are initially trained for 120 epochs, as described above. Replicating the transfer learning approach taken in [9], we then freeze the network up to and including the second fully connected layer on both sequence and epigenetics branch and train the remaining layers for a further 25 epochs. Dropout probability was 0.2 and the Adam learning rate was $10^{-3}$.

### Supplementary Figures

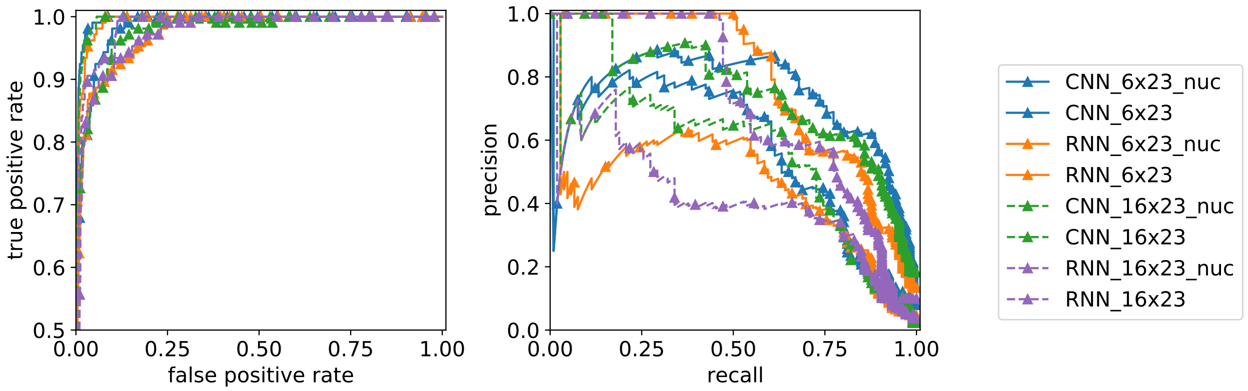


**Figure S1:** Receiver operating (left) and precision-recall (right) curves underlying Figure 3.

**Pairwise training** When combining various experimental studies, care must be taken as to their interaction during training. We therefore devised the notion of a pairwise training performance of two studies, which we define as the training performance when training on the larger and testing on the smaller of these studies. We did this for all studies in the crisprSQL dataset and visualised the result in a force-directed graph (Figure S2, Figure S3) where the force between nodes is proportional to the third power of the Spearman r obtained by pairwise training and testing.

Figure S2 shows that even though studies using the same cell lines show some bunching in a force-directed 2D graph where distance is proportional to generalisation from one study to the other, bunches heavily overlap between cell lines and experimental conditions. This supports the mixing of training data across studies despite differing experimental parameters. Experimental studies conducted on HEK293 cells appear to generalise best. Three distinct studies appear to generalise comparatively poorly to other studies, one of which was gained on HAP1 cells. Study [10] which has been gained on synthetic DNA appears to generalise to other studies about as well as studies on K562 cells.

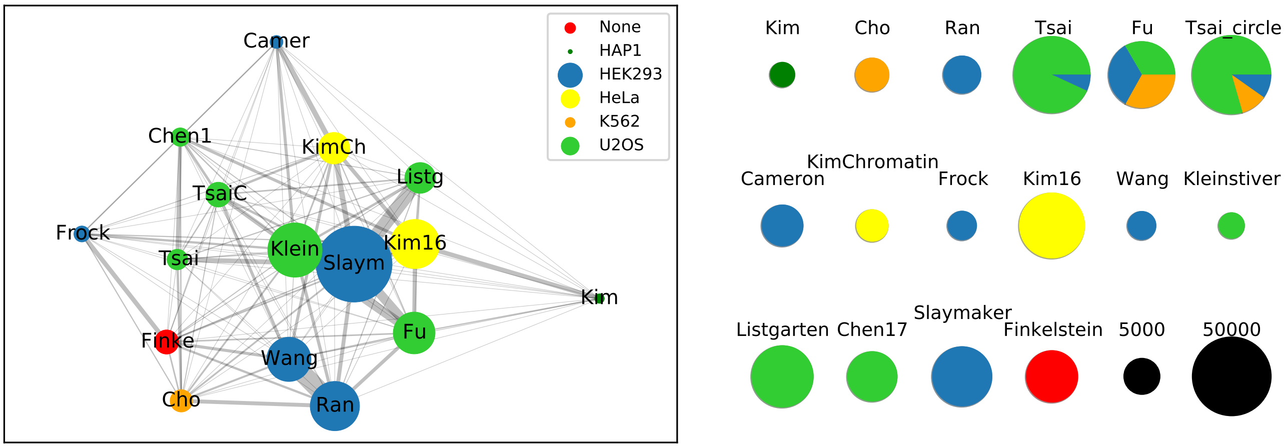


**Figure S2: Left panel**: 2D distance network representation of pairwise training performance using a Fruchterman-Reingold force-directed algorithm. The force between two nodes (studies) is proportional to the third power of the Spearman correlation when training on the larger and testing on the smaller of these studies. Study labels are abbreviated for better visibility. Edge width represents pairwise training performance of the two adjacent studies alone; bubble size indicates 'study importance', i.e. the overall summed performance of a study. Close positioning indicates good pairwise training performance. Studies have been coloured by majority cell line; all data has been gained using a CNN model which was shown to generalise fastest in terms of Spearman r in a separate experiment (data not shown). **Right panel**: Composition of the cell lines making up each individual study (colours as in left panel) with sizes proportional to the number of data points per study, including non-validated data points. Black circles act as a size legend.

Figure S7 shows that the RNN model is considerably influenced by the Nucleotide BDM feature which is calculated from the 147 base pair region around each nucleotide of the (off-)target DNA. This stands in

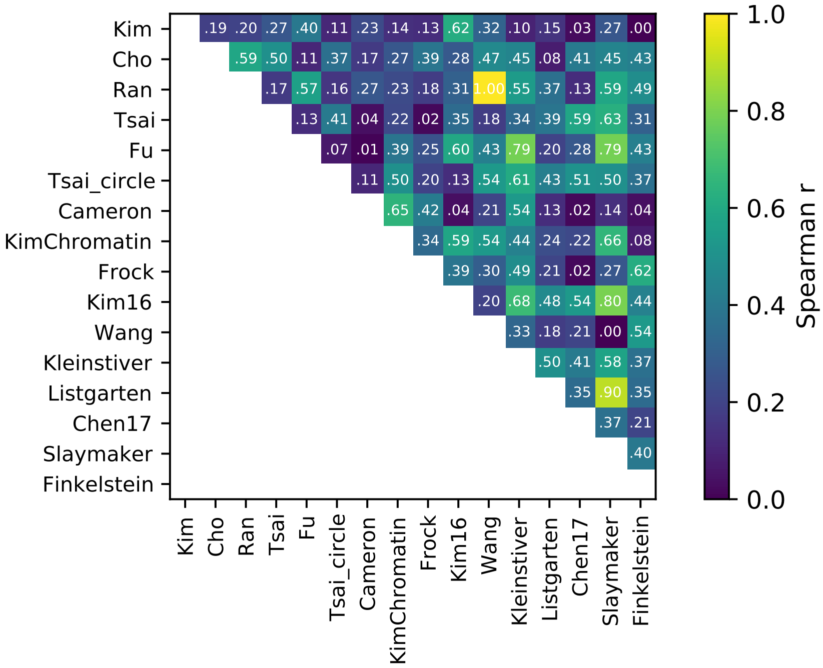


**Figure S3:** Spearman r data underlying Figure S2. Data has been gained using the CNN model.

contrast to the implicit assumption in most cleavage prediction models that sequence context only has a minor effect on cleavage. Our results emphasise the value of the said 147bp sequence context for prediction, since the feature has been shown to carry valuable information relevant to primary chromatin structure [7].

Sequence features pertaining to mismatched interfaces have a negative effect on cleavage activity for almost all interface types, supporting the notion from [14] that PAM-distal and PAM-proximal mismatches reduce cleavage activity due to cleavage inactivation and abrogation of DNA binding, respectively. We attribute the fact that this model gives little importance to energy features to the nature of recurrent neural networks penalising constant features across the recurrence dimension.

Figure S8 shows a strong dependence of the SHAP value for the $E_{\mathrm{CRISPRoff}}$ feature on the value of the feature itself. Small free energies of the DNA-RNA heteroduplex appear to favour cleavage activity, whilst very strong binding is not optimal for cleavage activity.

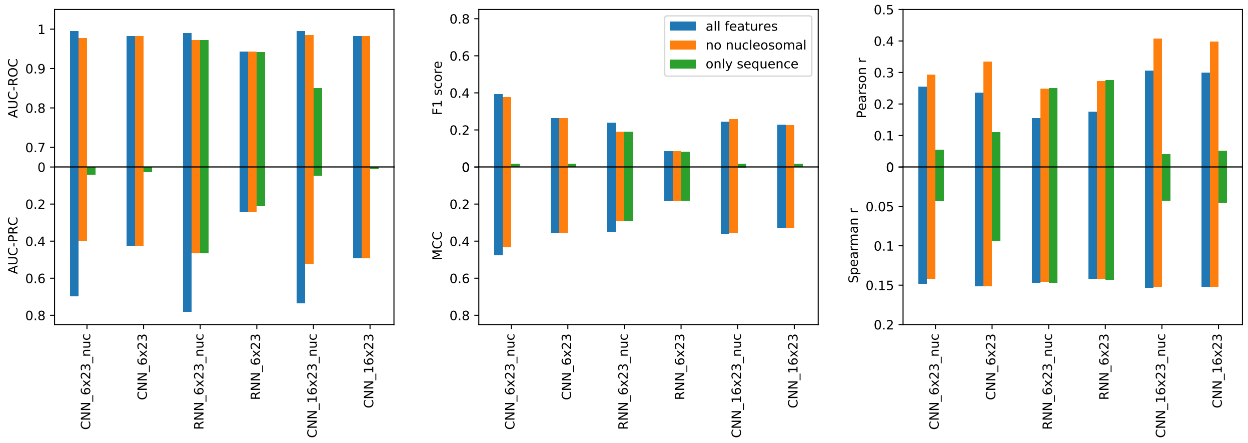


**Figure S4:** Performance comparison between piCRISPR models when tested on held out studies [11, 12, 13] whilst omitting certain sets of features. piCRISPR then uses default values for these features, leading to slightly reduced prediction accuracy.

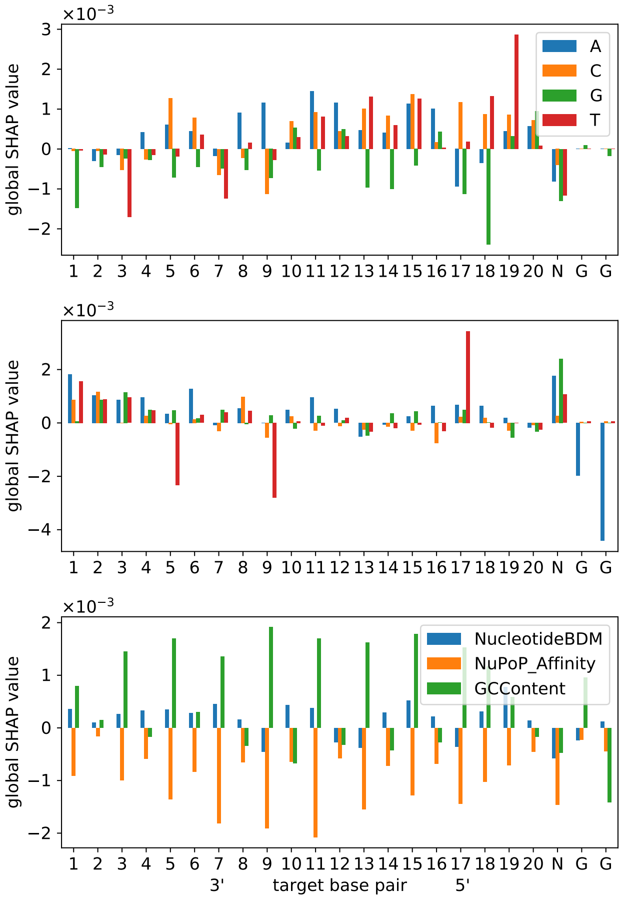


**Figure S5:** SHAP values for the $16 \times 23$ CNN classification model (Figure 4) in bar chart representation. The upper and middle bar plots refer to matched and mismatched interfaces, respectively. The different bar colours represent the respective nucleotides on the target protospacer DNA strand.

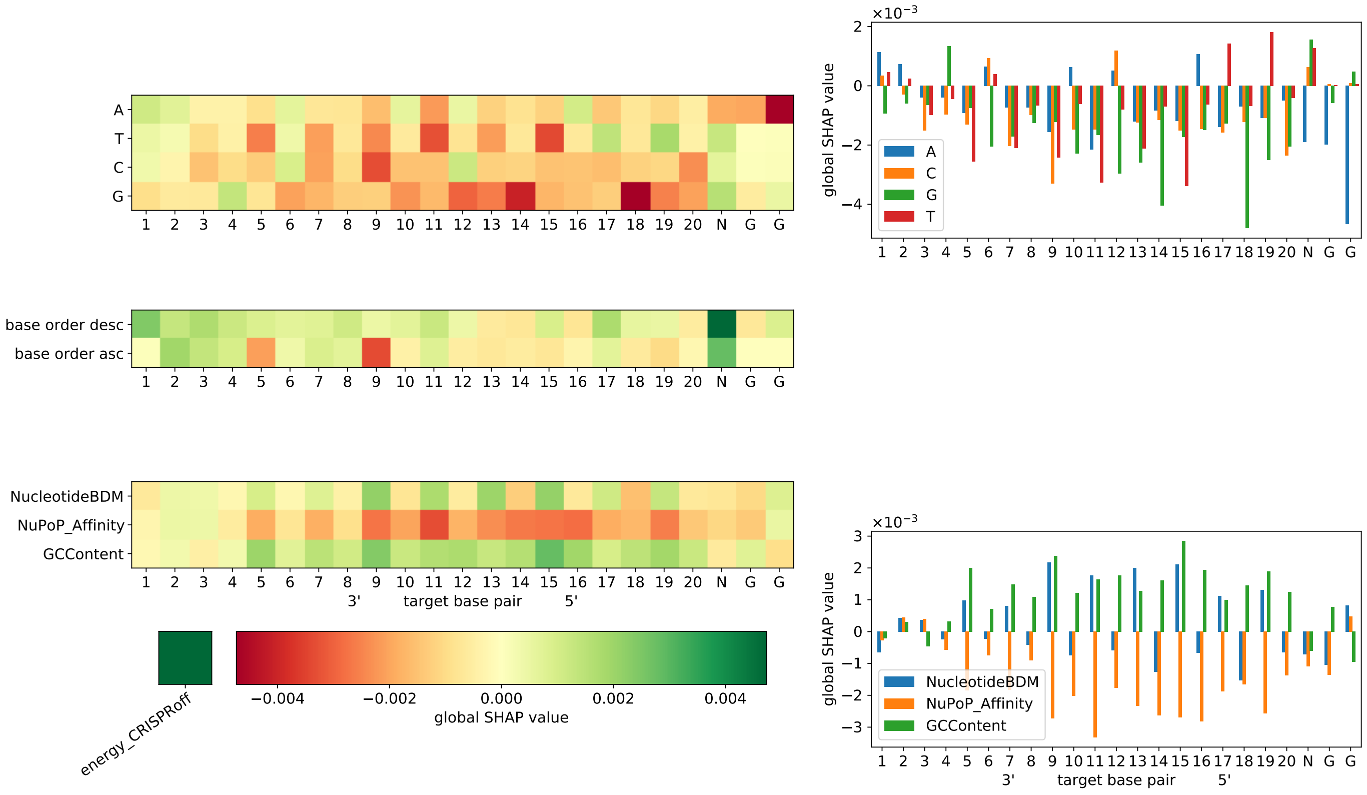


**Figure S6:** Base-pair resolved global SHAP values for the $6 \times 23$ CNN classification model. Training and test set are identical to Figure 4. The heatmap shows that after sequence features, the NuPoP Affinity channel contributes the most to the model prediction, with PAM-distal nucleotides less important than PAM-proximal ones.

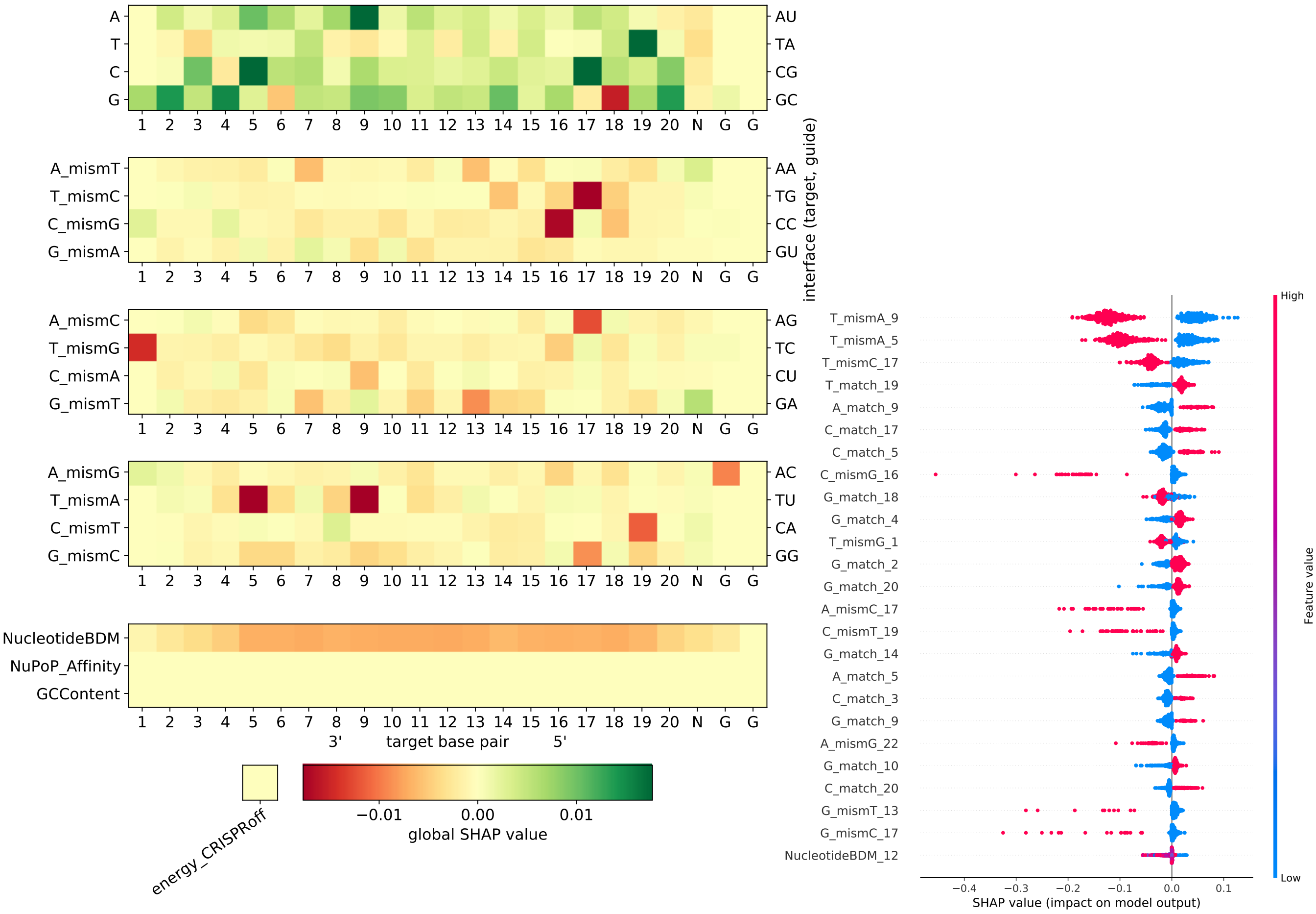


**Figure S7:** Base-pair resolved global SHAP values for the 16 × 23 RNN classification model. Training and test set are identical to Figure 4. The heatmap shows that after sequence features, the Nucleotide BDM channel contributes the most to the model prediction, with PAM-distal nucleotides less important than PAM-proximal ones.

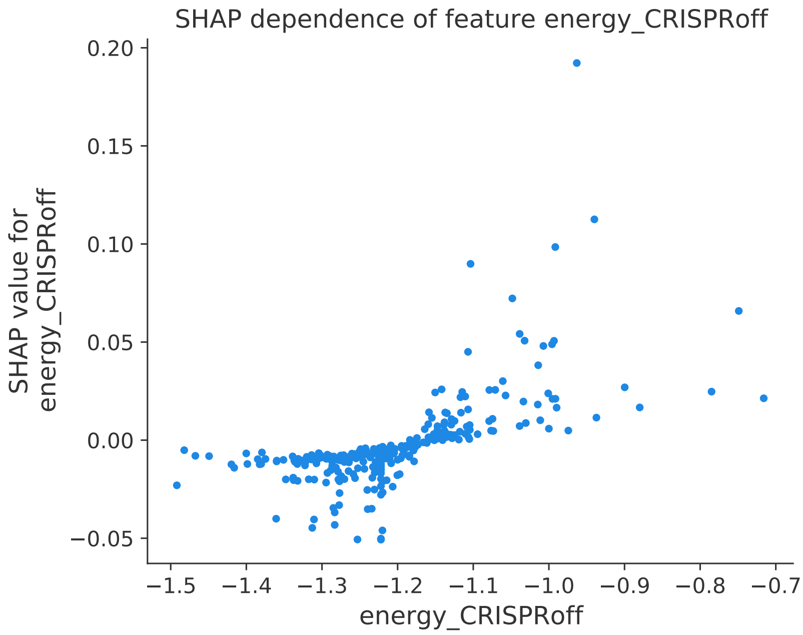


**Figure S8:** SHAP dependence plot for the $E_{\mathrm{CRISPRoff}}$ feature. The horizontal axis is in arbitrary units as provided by CRISPRoff [15]. Data has been gained using the 16 × 23 CNN classification model (Figure 4). For energy scores $E_{\mathrm{CRISPRoff}} > -1.15$, the RNA-DNA heteroduplex is weakly bound and opens easily, hence favouring cleavage (positive SHAP value), whereas for smaller $E_{\mathrm{CRISPRoff}}$ it becomes more tightly bound.

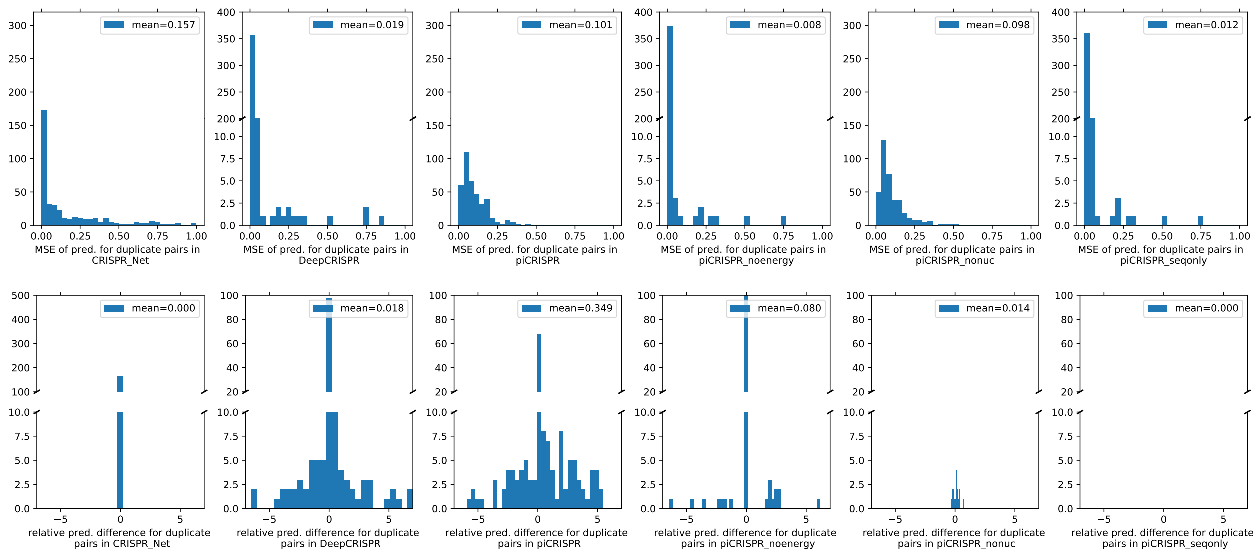


**Figure S9:** Distributions underlying Table 1.
